# Low oxygen post conditioning improves stroke-induced cognitive impairment

**DOI:** 10.1101/483453

**Authors:** Zidan Zhao, Lin Kooi Ong, Giovanni Pietrogrande, Sonia Sanchez Bezanilla, Kirby Warren, Marina Ilicic, Murielle G. Kluge, Clifford TeBay, Ole P. Ottersen, Sarah J Johnson, Michael Nilsson, Frederick R Walker

## Abstract

Post-stroke cognitive impairment has proven to be notoriously difficult to treat. In the current study, we sought to both better understand cellular changes that underpin cognitive deficits and to consider the potential restorative benefits of low oxygen post conditioning (LOPC). We were motivated to use LOPC as an intervention as it is one of the few experimental interventions previously shown to improve cognitive function post-stroke. Experimental stroke was induced by photothrombotic occlusion in adult male C57BL/6 mice. Mice were randomly assigned to either a normal atmospheric air exposure or low oxygen (11% O_2_) exposure groups three days post-occlusion. On day 17 post-stroke, mice were euthanized for histology or biochemical analyses. Stroked mice exposed to LOPC was associated with marked reductions in amyloid-beta both in its absolute level and in the extent of its oligomerization. Exposure to LOPC post-stroke also improved cellular deficits induced by stroke including an increase in vessel density, a reduction in vascular leakage, and restoration of AQP4 polarisation. Critically, stroked mice exposed to LOPC exhibited robust improvements in cognitive function post-stroke, assessed using a touchscreen based paired-associate learning task. These findings provide compelling pre-clinical evidence of the potential clinical utility of LOPC for enhancing recovery post-stroke.

## Introduction

While cognitive impairment is recognised to represent a major challenge for stroke survivors, there are currently no accepted clinical interventions for improving cognition post-stroke.^1, 2^ The paucity of effective pro-cognitive treatments has led to extensive pre-clinical efforts to develop innovative approaches to improve cognitive performance post-stroke.^3, 4^

One of the most promising pro-cognitive approaches identified to date has been intermittent exposure to a reduced oxygen environment.^5^ In the context of stroke, exposure to low oxygen environment prior to induction of an ischemic event has been shown to produce robust neuroprotection.^5^ More recently, the benefits of low oxygen exposure have been considered when deployed after an ischemic event, low oxygen post conditioning (LOPC), arguably a more translationally relevant timeframe to consider. We have recently identified that LOPC produces significant improvement in motor function and neuroprotection after experimental stroke.^6^ Further, LOPC has been demonstrated to enhance levels of neurogenesis and long-term memory function.^7^-^11^

Currently, it is not clear why cognitive deficits appear post-stroke nor why exposure to a low oxygen environment appears capable remediating these impairments. With respect to inducing cognitive deficits, several mechanisms are likely to be contributing factors, including loss of neural tissue and vasculature, vascular leakage, compromised expression of vascular receptors involved in metabolic waste removal as well as enhanced accumulation of neurotoxic amyloid-beta (Aβ) oligomers.^12, 13^ Increased levels of soluble Aβ accumulation, and in particular high molecular weight oligomers have been linked to cognitive decline.^14^

It has been shown that post-stroke exposure to LOPC promotes neuronal survival and vascular growth.^6, 15^ However, it unclear whether exposure to LOPC also improves other aspects of vascular function, or whether these improvements can modulate Aβ burden. As such, the current study sought to understand how stroke may impair key mechanisms that support cognition and how exposure to LOPC may influence these same mechanisms. To address these questions we exposed adult male mice to either regular atmospheric air post-stroke or to LOPC (11% oxygen, either 8h or 24h/day for 14 days). We assessed cognition using a rodent touchscreen platform for paired-associate learning (PAL) task, and investigated changes in cellular and molecular compositions of the peri-infarct territory.

## Materials and Methods

The data that support the findings of this study are available from the corresponding author upon reasonable request. A detailed materials and methods section is available in the Supplementary material.

### Ethical statements

Animal research was undertaken in accordance with ARRIVE guidelines. Experiments were approved by the University of Newcastle Animal Care and Ethics Committee (A-2013-338), and conducted in accordance with the New South Wales Animals Research Act and the Australian Code of Practice for the use of animals for scientific purposes.

### Animals

C57BL/6 male mice (eight weeks old) were obtained from the Animal Services Unit at the University of Newcastle. Mice were maintained in a temperature (21°C±1) and humidity-controlled environment with food and water available ad libitum. Lighting was on a 12:12 h reverse light–dark cycle (lights on 19:00 h) with all procedures conducted in the dark phase. Mice were habituated for a minimum of seven days prior to the start of the experiment.

### Randomization and Blinding

In all instances mice were randomly allocated to experimental groups. All outcome analyses were performed by independent study team members blinded to the treatment condition.

### Experimental design

A total of 128 C57BL/6 adult male mice were each randomly allocated to one of the following four groups: sham, stroke, stroke with exposure to 8h/daily of LOPC (LOPC 8h), and stroke with 24h/daily of LOPC (LOPC 24h). Each cohort of 32 mice (8 per group) were used for either behavioural testing, fixed tissue analysis (immunohistochemistry), western blotting, or PCR analysis. Brains and blood samples were collected at day 17 post-stroke.

### Animal surgery and low oxygen post conditioning

Photothrombotic vascular occlusion was performed as previously described.^16, 17^ Briefly, mice were anaesthetized by 2% isoflurane during surgical procedure on a temperature controlled (37°C ± 1) stereotaxic frame. The skull was exposed by incision of the skin along the midline of the scalp. Rose Bengal (200 μL, 10 mg/ml solution in sterile saline, Sigma-Aldrich, USA) was injected intraperitoneally. After 8 min, the skull was illuminated for 15 min by a 4.5 mm diameter cold light source positioned at 2.2 mm left lateral of Bregma, targeting the left motor and somatosensory cortices. For the sham group, the same surgical procedure was applied except Rose Bengal was replaced with 200 μL of sterile saline (0.9% NaCl, Pfizer, Australia). Mice were subjected to the low oxygen environment starting at 3 days post-stroke. Low oxygen exposure was achieved using a customized ventilated cage racking system that was retrofitted to accept 11% oxygen, provided by a pressure swing adaptor based hypoxic generator.^6^ The levels for CO_2_ were monitored and remained at atmospheric levels ∼350ppm and normal sea level atmospheric pressure (101kPa) within the conditioned chambers. Mice from the two exposure low oxygen conditions, LOPC 8h and LOPC 24h, were maintained in a low oxygen environment for either 8h (10am to 6pm) or 24h, respectively for two weeks. Sham and stroke only mice were handled for 2 min twice daily throughout the duration of experiment.

### Assessment of cognitive deficits

Associative memory cognitive domain was assessed in mice using the touchscreen platform for PAL task.^18, 19^ Touchscreen operant chambers (Campden Instruments Ltd., UK, Figure 1(a)) were used in the testing, and by their nature are inherently blinded.^18, 20^ A liquid reward (strawberry milkshake; Paul’s Milky Max) was provided to motivate animals’ performance. The task consists of two distinct phases: basic training, whereby the animal learns the association between making contact with the screen and the actual PAL task. Habituation/basic training was done before stroke and PAL task learning was commenced three days post-stroke. Eight animals per group (total 32 animals) were used in the PAL testing.

**Figure 1.**
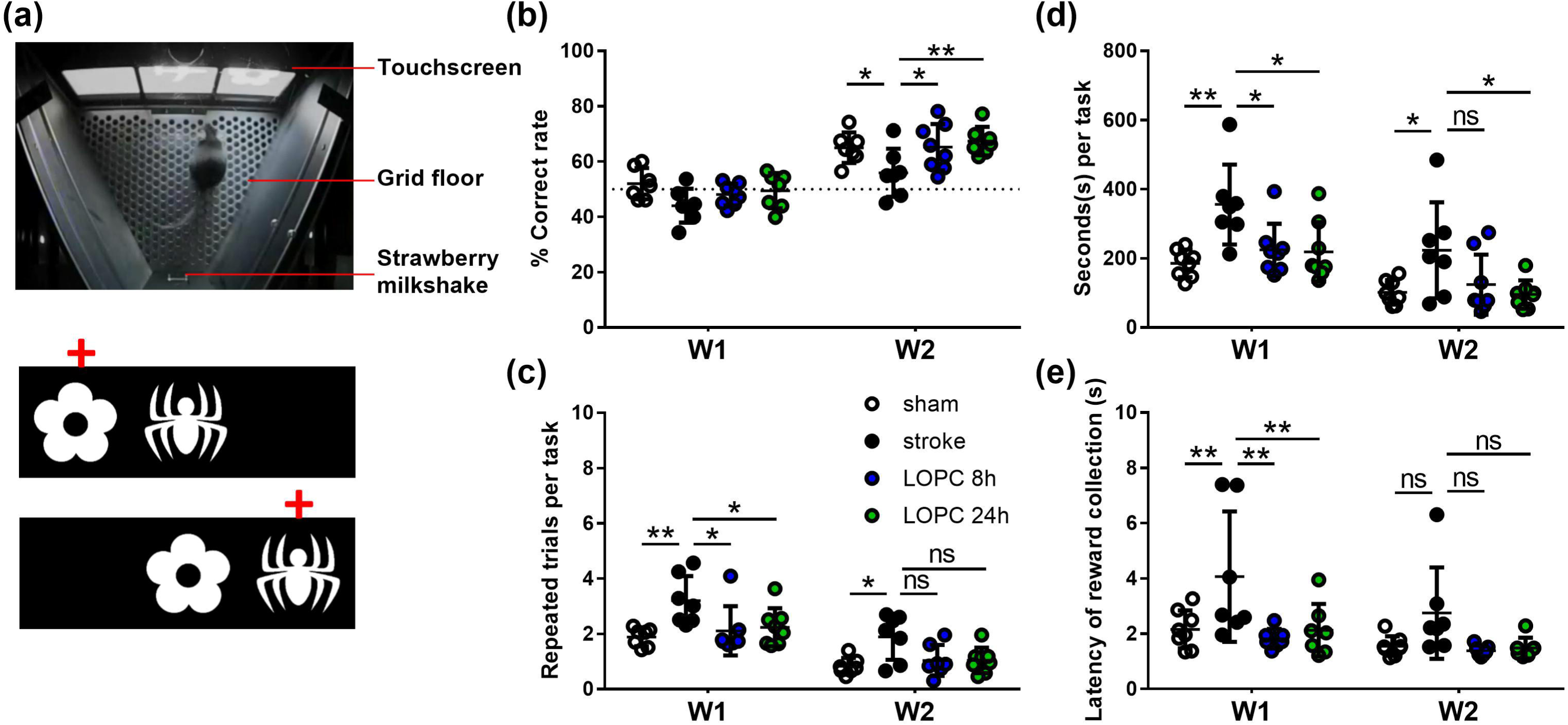
Illustration of the PAL task. **(a)** The Campden Instruments touchscreen chamber apparatus. To obtain the strawberry milkshake reward, the animal needs to choose the correct stimulus on the touchscreen. An illustration of the two different trial types and the correct location object pairing (red crosses) in PAL. Graphs show the animals’ performance of the four groups (sham, stroke, LOPC 8h and LOPC 24h) in **(b)** % correct rate, **(c)** repeated trials per task, **(d)** time per task and **(e)** latency of reward collection. Data expressed as mean±SD. ns: not significant, \**p*<0.05, \*\**p*<0.01 (two-way ANOVA, Tukey’s multiple comparisons).

#### Habituation/basic training

Before stroke, all mice were trained to touch the screen in order to receive the liquid reward. In order to receive a reward, animals were required to make contact with the screen when it was illuminated. Over 9 days all animals learnt to respond to screen illumination, with a minimum of 80% correct rate of response. Following general touchscreen training, mice underwent photothrombotic occlusion surgery.

#### PAL task learning

Three days post-surgery mice in the sham and stroke groups (±LOPC) commenced the PAL task. In the task, three stimuli images (a flower, plane and spider) were associated with a specific spatial location (left, center, right, respectively). In each trial, two images were displayed at the same time, one in the correct location and the other in an incorrect location (Figure. 1(a)). All trials were mouse initiated and independent of the experimenter. If the animal touched the image in its correct location, a reward was provided (a correct trial was recorded). After reward collection, the next trial was initiated. If the animal touched the incorrect image or the correct image in its incorrect location, it was punished by the absence of strawberry milkshake, no tone, and 5s house light on (incorrect trial). After 20s inter-trial interval, a repeat trial with presentation of the same stimuli was initiated (a repeated trial). This process was repeated until the mouse made the correct choice or 1h had elapsed. The number of repeated trials (also termed the perseveration index) was recorded and was not counted in the total trials administered or the correct rate (i.e % of correct trials). The time to finish each task was also recorded. The testing was terminated if the mouse successfully completed 36 trials or the testing session was 1 hour in length (which ever happened first).

### Haematocrit assessment

Blood haematocrit levels were measured using the i-STAT system and CG8 cartridges (Abbott Point of Care).

### Tissue processing

At day 17 post-stroke, mice were euthanized. For immunohistochemical analysis, animals (n = 8 per group) were deeply anesthetized via intraperitoneal injection of sodium pentobarbital and transcardially perfused with ice cold 0.9% saline for 2 mins followed by ice cold 4% paraformaldehyde (pH 7.4) for 13 mins. Brains were removed and post-fixed for 4 hours in the same fixative then transferred to a 12.5% sucrose solution in 0.1M PBS for storage and cyroprotection. Serial coronal sections were sliced on a freezing microtome (−25°C) at a thickness of 30μm. For western blot and PCR analysis, animals (n = 8 per group for western blot and PCR analysis, respectively) were deeply anesthetized via intraperitoneal injection of sodium pentobarbital and transcardially perfused with ice cold 0.1 % diethylpyrocarbonate in 0.9 % saline for 2 mins. Brains were dissected and rapidly frozen in −80°C isopentane. Sections were sliced using a cryostat (−20°C) at a thickness of 200μm and the peri-infarct territory (2 mm2 around infarct core) were punched using a 1 mm tissue punch. Samples were kept frozen at all times until protein and mRNA extraction.

### Histology and immunohistochemistry analysis

For immunoperoxidase labelling, free floating sections were immunostained as previously described^17, 21^, with one of the following primary antibodies: mouse anti-NeuN, mouse anti-GFAP, rabbit anti-Iba-1, rabbit AQP4, biotinylated goat anti-IgG. All antibodies details have been provided in Table I. Sections to be immunolabelled with rabbit anti-collagen IV additionally underwent pepsin antigen retrieval using the method described by S. Franciosi et al.^22^ Sections were rinsed with 0.1 M PB and endogenous peroxidases were quenched in 0.1 M PB containing 3% hydrogen peroxide. Non-specific binding was blocked with 3% normal horse serum. Sections were incubated in primary antibody with 2% normal horse serum for 48 hours at 4 C° and then were washed in 0.1 M PBS for 30 min and incubated with a biotinylated secondary antibody of corresponding species for 2h at room temperature, rinsed, incubated in 0.1% extravadin peroxidase for 1h, and then rinsed again. Immunolabelling was developed using a nickel-enhanced 3, 3’-diaminobenzidine (DAB) reaction. Tissues from the four experimental groups were performed simultaneously and the DAB reactions were developed for exactly the same length of time following the addition of glucose oxidase (1:1000). Negative control sections, in which no primary antibodies were added, were developed at the same time to confirm the specificity of labelling. After processing was completed sections were washed, mounted onto chrome alum-coated slides and cover-slipped.

**Table I.** List of antibodies used for western blot and immunohistochemistry.

| Target | Sources of antibodies | Application | Dilution |
| --- | --- | --- | --- |
| **Collagen IV** | Abcam, rabbit anti-collagen IV, ab6586 | IHC | 1:1000 |
| **GFAP** | Sigma, mouse anti-GFAP, G3893 | IHC | 1:1500 |
| **Iba-1** | WAKO, rabbit anti-Iba-1, 019-19741 | IHC | 1:1000 |
| **NeuN** | Millipore, mouse anti-NeuN, MAB377 | IHC | 1:500 |
| **AQP4** | Millipore, rabbit anti- AQP4, AB3594 | IHC | 1:1000 |
| **IgG** | Abcam, biotinylated goat anti-mouse IgG, ab64255 | IHC | 1:500 |
| **amyloid $\beta$** | Cell Signaling Technology, anti-amyloid $\beta$ (D3D2N), #15126 | WB | 1:1000 |
| **β-actin** | Sigma-Aldrich, Monoclonal anti-β-actin-HRP, A3854 | WB | 1:50000 |
| **Rabbit IgG** | Jackson ImmunoResearch, goat anti-rabbit-biotin, #111-065-003 | IHC | 1:500 |
| **Mouse IgG** | Biorad, anti-Mouse-HRP antibody, #170-6516 | WB | 1:10000 |
|  | Jackson ImmunoResearch, goat anti-mouse-biotin, #115-065-003 | IHC | 1:500 |

WB, western blot; IHC, immunohistochemistry.

### Image acquisition, tissue loss, cell count, thresholding and AQP4 polarization analysis

Images were acquired at 20 X using Aperio AT2 (Leica, Germany). ImageJ (1.50, NIH) and Matlab (R2015a, MathWorks) were used to quantitate tissue loss, cell count, Aquaporin-4 (APQ4) vessel/parenchymal ratio, and to analyse intensity and area coverage of immunolabelling.^16, 17, 19, 23-25^ To estimate tissue loss within the infarcted hemisphere, the area of contralateral and ipsilateral hemispheres were measured across four sections (at Bregma level +1.0, 0.0, −1.0 and −2.0 mm, Figure 2(a)) using Image J. The % tissue loss was determined by the equation: [(average area of contralateral hemisphere – average area of ipsilateral hemisphere)/ area of contralateral hemisphere] x 100 %. The quantitative analysis was undertaken specifically in the peri-infarct territory as defined by 0.01mm from infarct, the region was 0.25mm by 0.5mm in size. Cumulative threshold analysis was performed using Matlab functions. Firstly, for each of the acquired images, the number of pixels occurring at each of the pixel intensities was determined. The pixel intensity values are then rank ordered 0-255 along with the corresponding number of pixel that occur at each value. For the purposes of analysis, we calculated the percentage of cumulative threshold material for the range of pixel intensity values (Figure 7). Pixel intensity level considered to be optimal for detecting genuine differences in immunoreactive signal was determined using ImageJ software to visualize thresholding of cropped regions at individual pixel intensities. This threshold level was used to investigate group differences for all labels. For NeuN, GFAP and Iba-1 positive cell counts, exhaustive manual cell counts were undertaken within the cropped regions (Bregma 0.0mm, Figure 2(a)). The vessel digital reconstruction was performed as previously described.^19, 26, 27^ Collagen IV positive cells were isolated from the background using multi-level Otsu’s thresholding method, which calculates the threshold that minimizes the interclass pixel intensity variance between groups. Using Matlab functions we determined percentage area covered by the labelling. AQP4 analysis involved quantifying the intensity of AQP4 labelling on the vessel lumen relative to that in the adjacent parenchyma (APQ4 vessel/parenchymal ratio).

### Biochemical analysis

Protein homogenates from peri-infarct samples were obtained and Western blotting was performed as previously described.^16, 17^ PVDF membranes were incubated with primary antibody anti-amyloid β (D3D2N) overnight at 4°C, followed by secondary antibody anti-Mouse-HRP antibody for 1h at 25°C. In between each incubation step, membranes were washed in TBS-T. Membranes were visualized on Amersham Imager 600 using Luminata Forte Western blotting detection reagents. The density of the bands was measured using Amersham Imager 600 Analysis Software. RNA was isolated from the peri-infarct samples using the Illustra RNAspin Kit (GE Healthcare, Cat#25-0500-70) according to manufacturer’s specifications. Quantitative RT-PCR was performed as previously described.^28^ cDNA was generated using SuperScript™ III First Strand Synthesis System for RT-PCR (Invitrogen, Cat# 18080-044) according to manufacturer’s instructions on a GeneAmp PCR System 9700 instrument (Applied Biosystems). Quantitative RT-PCR was performed on an Applied Biosystems® 7500 (Applied Biosystems) or ViiA7 (Thermofisher) instruments using SensiFAST SYBR® Lo-ROX Master Mix (Bioline, Cat# BIO-94020). Genes of interest (Table II) were normalized on the housekeeping gene GAPDH and data are expressed as 2-ΔΔCt as fold change relative to sham.

**Table II.**
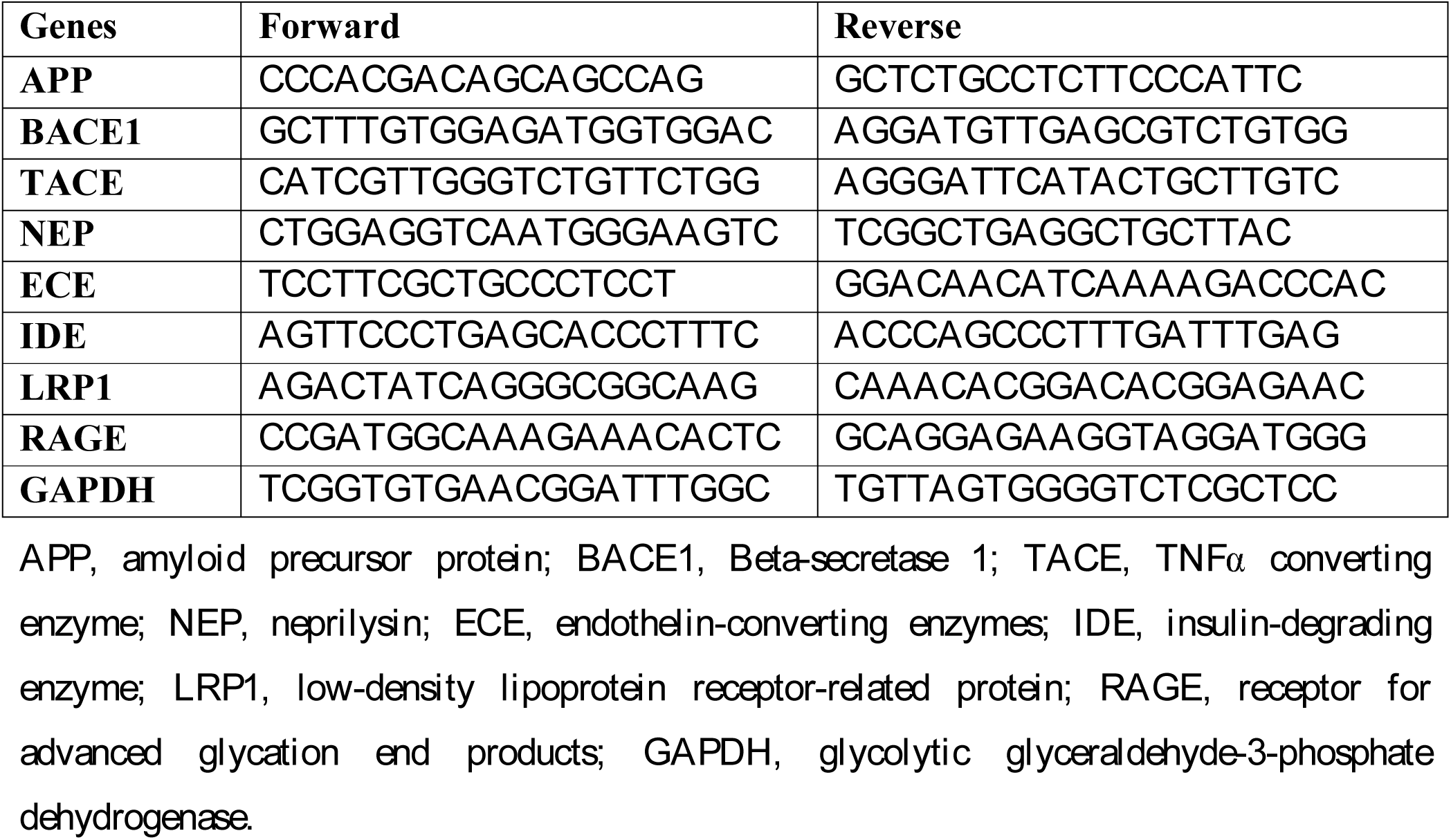
PCR primer sequences.

**Figure 2.**
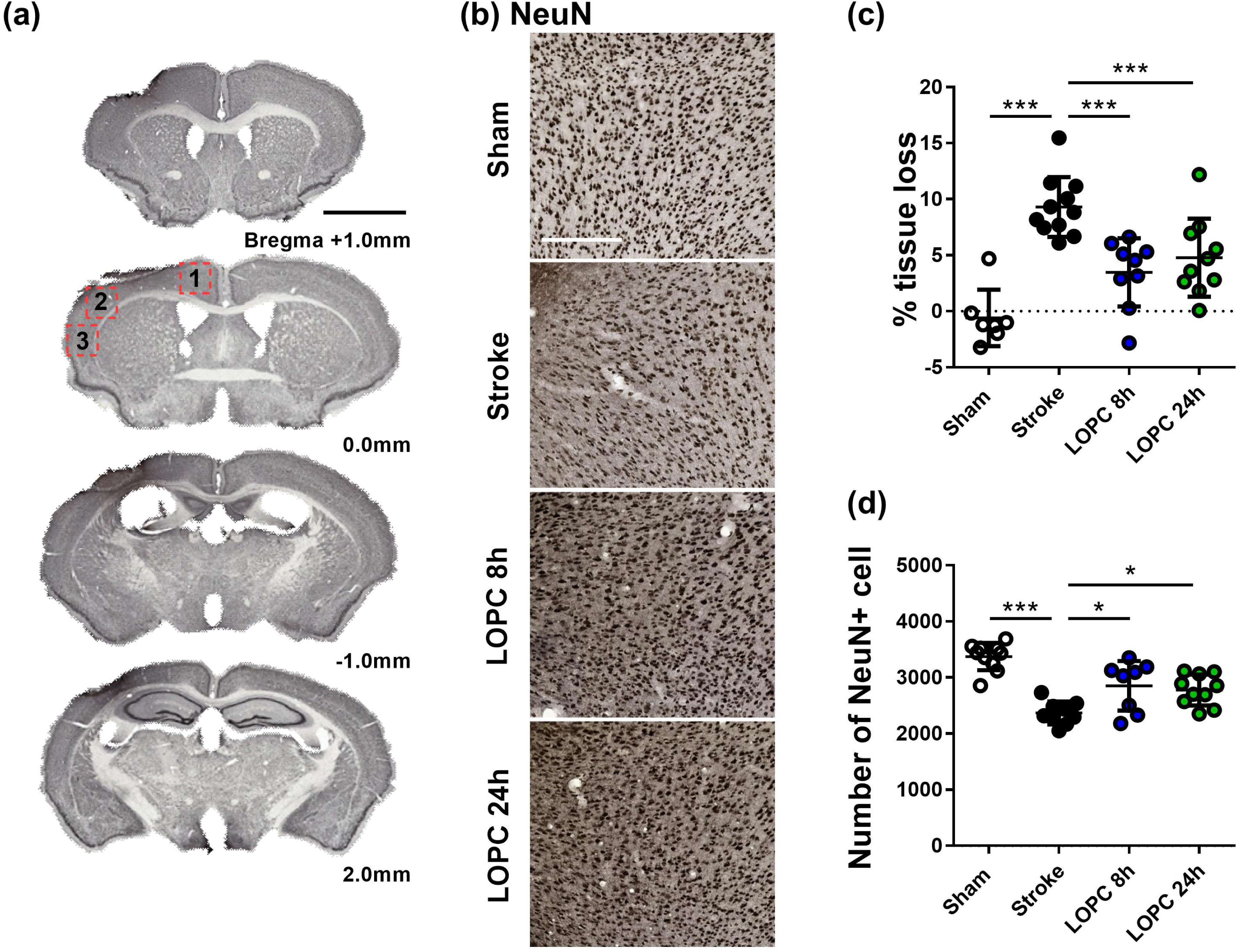
Illustration of the neural tissue loss. **(a)** The stroke sections from Bregma +1.0mm to Bregma −2.0mm. **(b)** Representative labelling for NeuN for the four groups: sham, stroke, LOPC 8h and LOPC 24h. **(c)** The graph shows that LOPC 8h and LOPC 24h animals had significant lower % tissue loss compared to the stroke only animals. **(d)** The graph illustrates the total number of NeuN positive cells for four groups. Multiple peri-infarct regions (red boxes) at Bregma 0.0 were included for neuronal cell counts. Data expressed as mean±SD. \*\*\**p*<0.001 (one-way ANOVA, Tukey’s multiple comparisons). White scale bar represents 300μm and black scale bar represents 1mm.

### Statistical analysis

All data for sham, stroke, LOPC 8h and LOPC 24h groups were expressed as mean±SD and were analysed using Prism 6 for Windows Version 6.01, GraphPad Software. Two-way ANOVA was used to determine whether there were time and treatment effects across groups in the PAL task. All other experiments used One-Way ANOVA to determine whether there were any significant treatment effects across the groups. Additional Tukey multiple comparisons were used to analyse differences between the mean of each group and the mean of every other group. The significant differences shown on the graphs with asterisks (*) refer to the post hoc tests. All differences were considered to be significant at *p*<0.05.

## Results

### Low oxygen post conditioning increases haematocrit in mice

To confirm the biological effect of the LOPC protocol, haematocrit assessment was performed 14 days following LOPC treatment (17 days post-stroke). We identified that there was no significant difference between sham and stroke animals (0.38±0.006 vs. 0.38±0.005, *p*=0.97). The stroked LOPC 8h (0.48±0.005) and LOPC 24h (0.49±0.008) animals had a significantly higher haematocrit than stroke only animals (*p*<0.001 and *p*<0.001, respectively).

### Low oxygen post conditioning ameliorates cognitive deficit after stroke

In order to assess whether LOPC treatment improved cognitive outcomes post-stroke, animals were exposed to LOPC for 8h or 24h alongside daily PAL touchscreen task for 2 weeks (Figure 1(a)).

Analysis of the *% correct responses* indicated that there was a main effect for each of treatment group (F=118.3, p<0.001) and time (F=4.16, *p*<0.05) (Figure 1(b)). In the second week of testing we identified that the animals exposed to stroke performed significantly lower than the sham (*p*<0.05), LOPC 8h (*p*<0.05) and LOPC 24h (*p*<0.01) groups.

Analysis of the *number of repeated trials per task* revealed a significant main effect for group (F=77.90, *p*<0.001) and time (F=6.36, *p*<0.01) (Figure 1(c)). In the first week, stroked animals performed significantly more repeated trials when compared to the sham (*p*<0.01), LOPC 8h (*p*<0.05) and LOPC 24h (*p*<0.05) groups. In the second week of training stroked animals still performed a greater number of repeated trials when compared to sham (*p*<0.05), however there was no significant difference between stroke and LOPC 8h (*p*=0.08) or LOPC 24h animals (*p*=0.07).

Analysis of *seconds per task* revealed a significant main effect for group (F=5.60, *p*<0.01) and time (F=84.59, *p*<0.001) (Figure 1(d)). In the first week, stroked animals took significantly longer time per task when compared to the sham (*p*<0.01), LOPC 8h (*p*<0.05) and LOPC 24h (*p*<0.05) groups. In the second week of training stroked animals still took significantly longer time per task when compared to sham (*p*<0.05) and LOPC 24h (*p*<0.05) animals, however there was no significant difference between stroke and LOPC 8h animals (*p*=0.11).

Analysis of *latency of reward collection* revealed a significant main effect for group (F=5.27, *p*<0.01) and time (F=14.03, *p*<0.001) (Figure 1(e)). In the first week, stroked animals took significantly longer time to collect reward when compared to the sham (*p*<0.01), LOPC 8h (*p*<0.01) and LOPC 24h (*p*<0.01) groups. There was no statistically significant deference between groups in the second week.

[insert Figure 1.]

### Low oxygen post conditioning reduces tissue loss and neuron loss after stroke

Total brain tissue volume and neuronal cell population were examined using immunohistochemistry (Figure 2). There was a statistically significant increase in average volume of tissue loss in stroke animals compared to sham animals (*p*<0.001). The average volume of tissue loss in the LOPC 8h and LOPC 24h animals was significantly smaller than observed in stroke only animals (F=16.68, *p*<0.001 and *p*<0.001, respectively, Figure 2(c)). Mature neuronal marker NeuN was used to assess the neuronal cell loss in the peri-infarct regions after stroke. Exhaustive manual cell counts were undertaken within the cropped regions indicated in Figure 2(a). Stroke animals significantly reduced numbers of NeuN positive cells in the peri-infarct territory compared to sham animals (F=18.44, *p*<0.001), a state that was restored in both LOPC 8h and LOPC 24h animals (*p*<0.05 and *p*<0.05, respectively. Figure 2(d)).

[insert Figure 2.]

### Low oxygen post conditioning promotes cerebrovascular remodelling and microglia activation

Cerebrovascular remodelling was examined by threshold analysis and digital vessel reconstruction of Collagen IV labelling in order to determine whether LOPC stimulated angiogenesis (Figure 3(a)). For thresholding analysis, the data for each group was expressed as a fold increase of the mean±SD relative to the mean of the sham group (for cumulative threshold analysis see the Supplementary material). Stroke animals had significantly reduced Collagen IV immunoreactivity levels and percentage area covered by Collagen IV positive cells in the peri-infarct territory, compared to sham animals (F=28.47, *p*<0.05 and F=10.02, *p*<0.05, respectively). Both LOPC 8h and LOPC 24h significantly promoted the density and distribution of Collagen IV compared to the stroke only animals.

**Figure 3.**
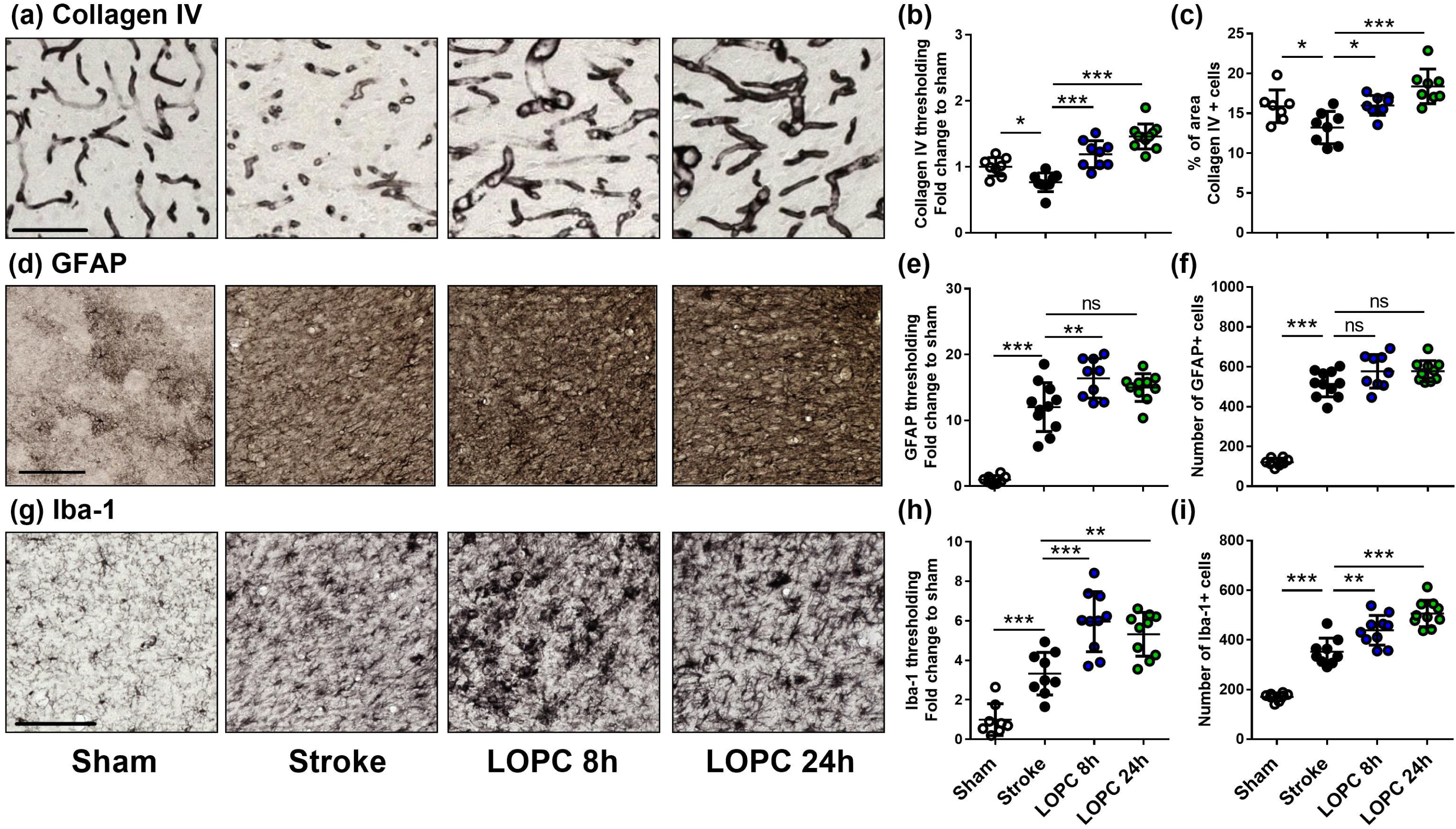
The effects of LOPC on vascular and glial cells within the peri-infarct regions. Four left panels in each row illustrate representative labelling for each marker investigated **(a)** Collagen IV, **(d)** GFAP and **(g)** Iba-1 for the four groups: sham, stroke, LOPC 8h and LOPC 24h. The first set of graphs **(b, e, h)** show quantification of the fold change of thresholded material for each of the markers. The second set of graphs show **(c)** percentage of area covered by Collagen IV positive cells, and number of **(f)** GFAP positive cells and **(i)** Iba-1 positive cells. Data expressed as a fold change of mean±SD for each group relative to the mean of the sham group. For cumulative threshold analysis refer to the Supplementary material. \**p*< 0.05, \*\*\**p*<0.001 (ANOVA, Tukey’s multiple comparisons). All scale bars represent 100μm.

To investigate whether LOPC influenced glial cells in the peri-infarct territory following stroke, we performed threshold analysis and cell counts using astrocyte marker, GFAP, and microglia marker, Iba-1 (Figure 3(d) and (g)). Stroke induced a significant increase in both GFAP and Iba-1 immunoreactivity relative to sham animals (F=55.06, *p*<0.001 and F=31.72, *p*<0.001, respectively). This corresponded with an increase in the number of GFAP and Iba-1 positive cells in stroke animals compared to sham animals (F=108.5, *p*<0.001 and F=72.68, *p*<0.001, respectively). LOPC 8h exhibited modestly elevated thresholded immunoreactivity levels of GFAP to stroked animals. LOPC 8h and LOPC 24h significantly increased thresholded immunoreactivity levels and Iba-1 positive cells to stroke animals.

[insert Figure 3.]

### Low oxygen post conditioning reduces Aβ oligomer accumulation after stroke

The tissue samples from the peri-infarct territory were analysed by western blot using anti-Aβ for soluble Aβ oligomers. The results for Aβ oligomers at 56kDa (dodecamer), 50kDa (decamer), 25kDa (pentamer), 5kDa (monomer) and total Aβ (5-200kDa) levels were normalized to β-actin as loading control (Figure 4). Data for all groups were expressed as a fold increase of the mean±SD for each group relative to the mean of the sham group. All Aβ oligomers showed similar patterns. Specifically, at 56kDa, 50kDa, 25kDa, and total Aβ levels, were elevated in the stroke group relative to sham animals (F=10.44, *p*<0.001; F=28.60, *p*<0.001; F=12.10, *p*<0.001 and F=12.14, *p*<0.001, respectively). Both the LOPC 8h and LOPC 24h displayed lower levels of oligomerization relative to the stroke alone condition (LOPC 8h; *p*<0.05, *p*<0.001, *p*<0.05, *p*<0.05 and LOPC 24h; *p*<0.01, *p*<0.001, *p*<0.05, *p*<0.001, respectively). At 5kDa level, stroke induced a significant increase relative to sham animals (F=3.01, *p*<0.05), but there was no significant difference compared to LOPC 8h and LOPC 24h (*p*=0.23 and *p*=0.15, respectively).

**Figure 4.**
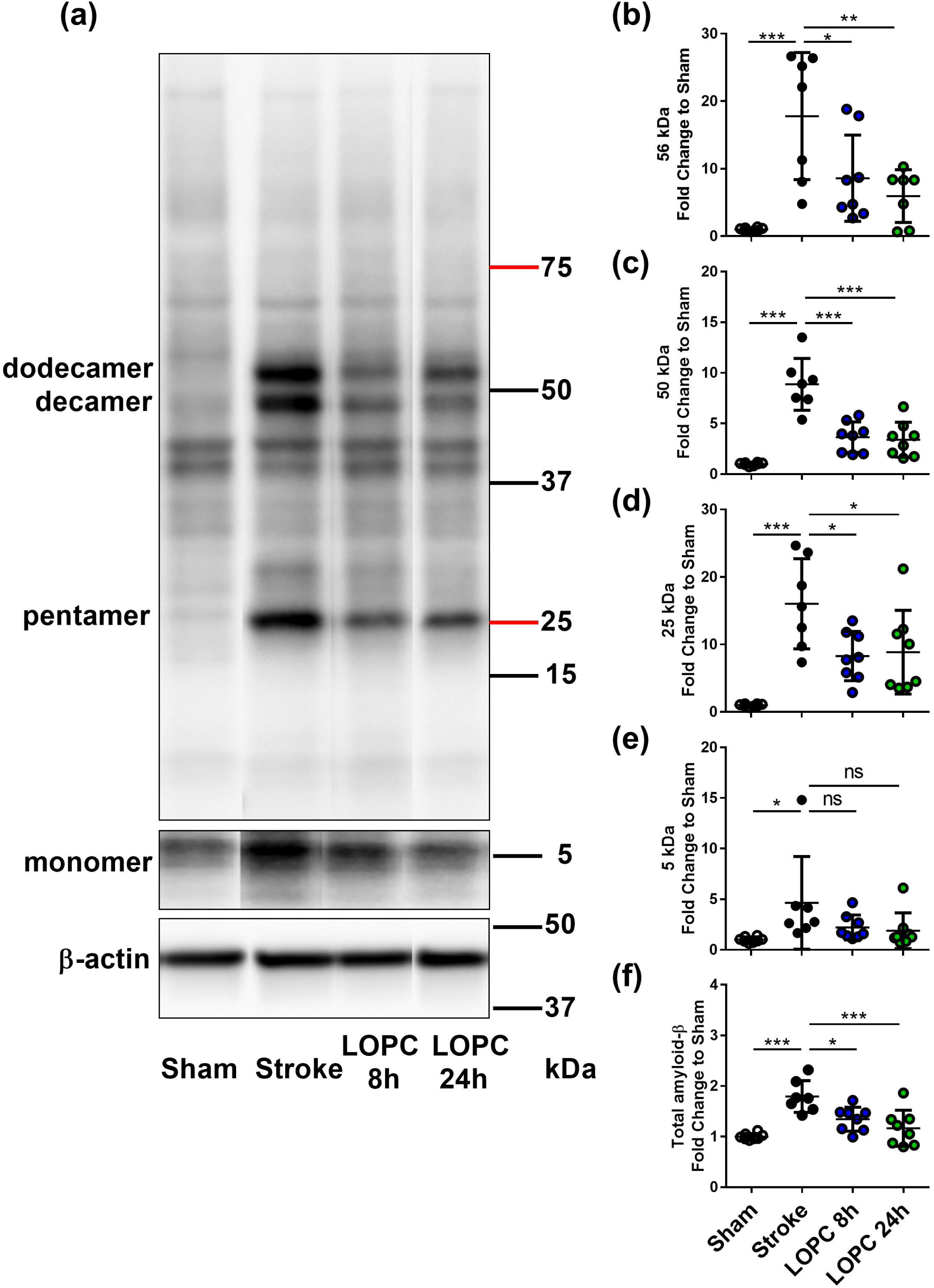
LOPC reduces Aβ in the peri-infarct territory after stroke. **(a)** The left panel is the representative western blot of protein samples in peri-infarct territory from sham, stroke, LOPC 8h and LOPC 24h animals. Bands were detected using D3D2N anti-Aβ antibody. Loading controls were performed by analysis of β-actin. The bar graphs at the right are quantification of Aβ oligomers at **(b)** 56kDa (dodecamer), **(c)** 50kDa (decamer), **(d)** 25kDa (pentamer), **(e)** 5kDa (monomer) and **(f)** total Aβ (5-200kDa) deposition. Data expressed as a fold change of mean±SD for each group relative to the mean of the sham group. ns: not significant, \**p*<0.05, \*\**p*<0.01 \*\*\**p*<0.001 (one-way ANOVA, Tukey’s multiple comparisons).

[insert Figure 4.]

### LOPC alters APP and BACE mRNA expression

We investigated changes in the expression levels of several key genes involved in the generation, degradation and export of Aβ using qPCR (Figure 5), specifically amyloid precursor protein (APP); beta-secretase (BACE); TNFα converting enzyme (TACE); neprilysin, (NEP); endothelin-converting enzyme (ECE); insulin-degrading enzyme (IDE); low-density lipoprotein receptor–related protein-1 (LRP1); and receptor for advanced glycation end products (RAGE). Stroke animals exhibited a significant decrease in the expression of APP and BACE mRNA levels relative to sham animals (F=16.56, *p*<0.001 and F=4.61, *p*<0.05, respectively). This reduction of APP and BACE mRNA levels was reversed by both LOPC 8h and LOPC 24h. Stroke alone induced a significant increase in the expression of TACE and NEP mRNA levels relative to sham animals (F=11.13, *p*<0.001 and F=20.62, *p*<0.001). However, stroke alone induced a significant decrease in the expression of ECE mRNA levels relative to sham animals (F=17.14, *p*<0.001). LOPC 8h modestly elevated LRP1 and RAGE mRNA levels (*p*<0.001 and *p*<0.05, respectively), and LOPC 24h increased ECE mRNA levels ((*p*<0.001), relative to stroke only animals.

**Figure 5.**
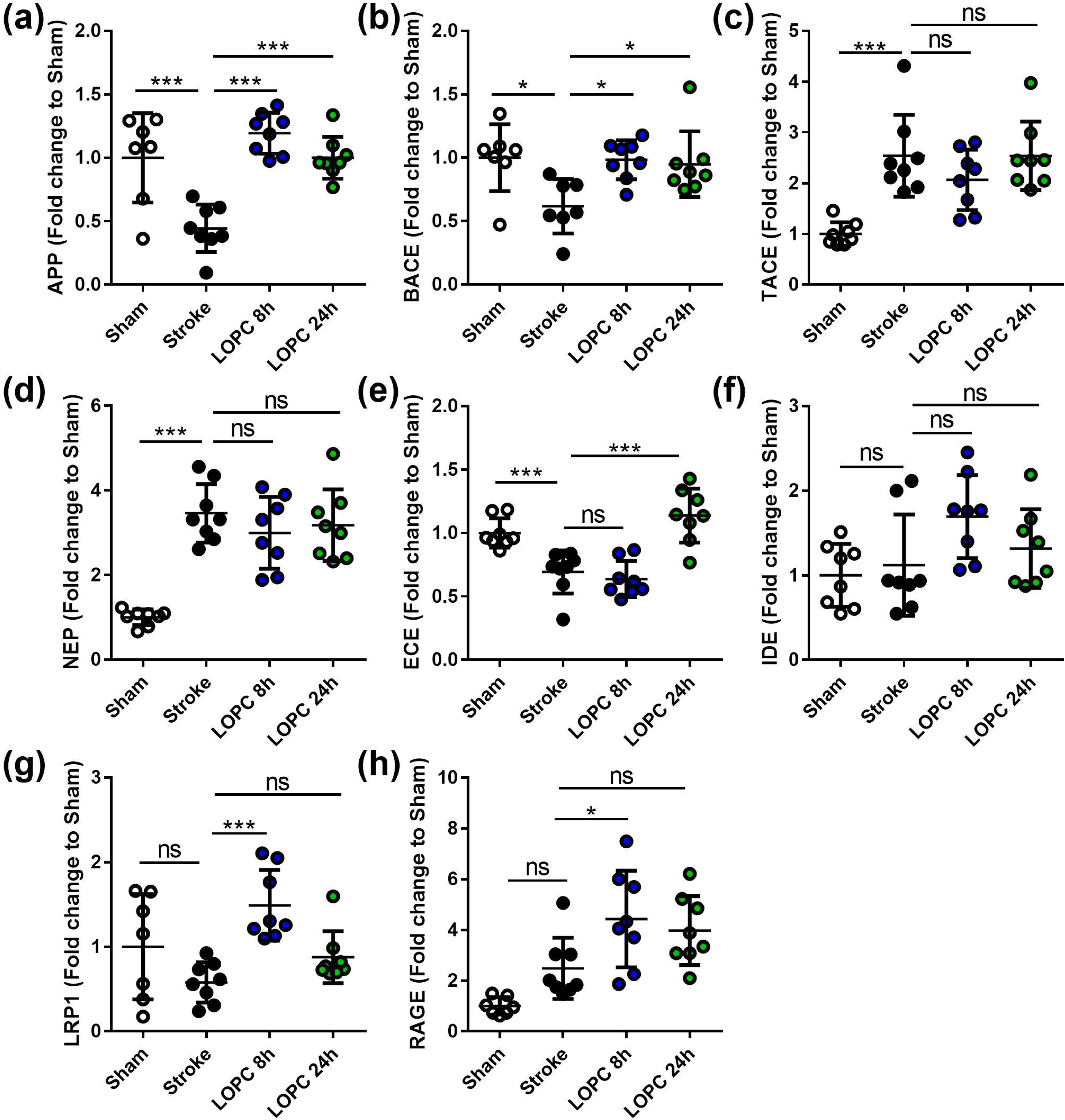
The expression levels of genes involved in the generation, degradation and export of Aβ. The expression of **(a)** amyloid precursor protein, APP; **(b)** beta-secretase, BACE; **(c)** TNFα converting enzyme, TACE; **(d)** neprilysin, NEP; **(e)** endothelin-converting enzyme, ECE; **(f)** insulin-degrading enzyme, IDE; **(g)** low-density lipoprotein receptor– related protein-1, LRP1; and **(h)** receptor for advanced glycation end products, RAGE. Data expressed as a fold change of mean±SD for each group relative to the mean of the sham group. ns: not significant, \**p*<0.05, \*\*\**p*<0.001 (one-way ANOVA, Tukey’s multiple comparisons).

[insert Figure 5.]

### LOPC improves vascular leakage and AQP4 polarization after stroke

Stroke-induced cerebrovascular leakage was assessed by IgG staining in the peri-infarct regions (Figure 6(a)). A significant increase in IgG (F=13.87, *p*<0.001) was present in stroke animals compared to sham animals. Vascular leakage was improved significantly by exposure to LOPC 24h (*p*<0.001), however there was no significant difference between LOPC 8h and stroke only animals (*p*=0.45).

**Figure 6.**
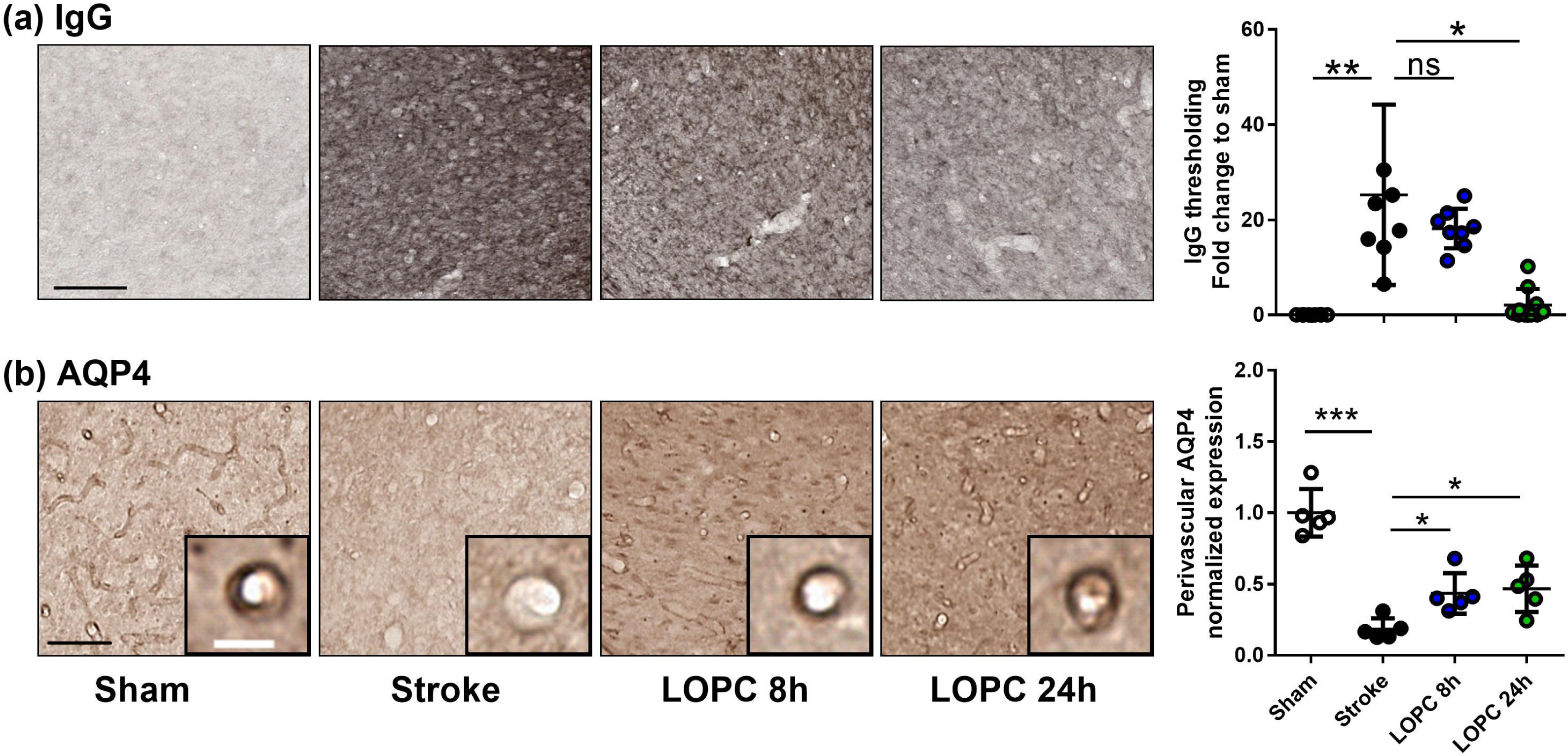
LOPC improves vascular leakage and AQP4 polarity within peri-infarct region following stroke. **(a)** Images illustrate representative labelling of IgG staining, an index of cerebrovascular leakage. **(b)** Images illustrate representative labelling of AQP4. Insets show APQ4 polarity on vessels at high magnification. The right bar graph illustrates the AQP4 polarization. Data expressed as a fold change of mean±SD for each group relative to the mean of the sham group. ns: not significant, \**p*<0.05, \*\**p*<0.01 \*\*\**p*<0.001 (one-way ANOVA, Tukey’s multiple comparisons). Black scale bar represents 100μm and white scale bar represents of inset represents 10μm.

**Figure 7.**
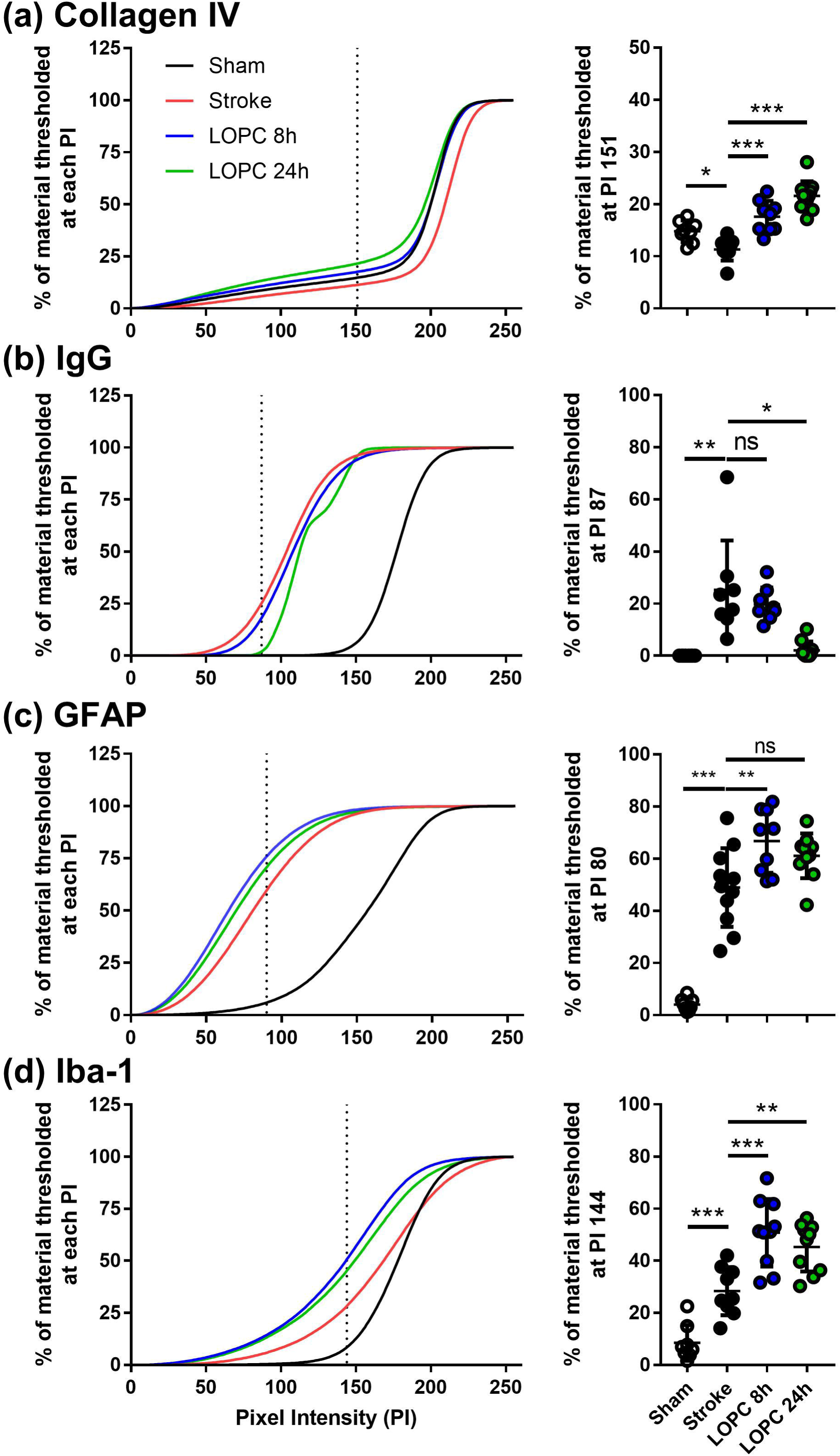
Cumulative threshold analysis. Left panels show the percentage of material thresholded of **(a)** Collagen IV, **(b)** IgG, **(c)** GFAP and **(d)** Iba-1 immunolabels at different levels of pixel intensity (PI). Dotted line are PI level considered to be optimal for detecting genuine differences in immunoreactive signal. The right most panel illustrates quantification of the change % of thresholded material for each of the immunolabels at selected PI. Data for left panels are presented as mean and data for right panels are presented as mean±SD. *p<0.05, ***p<0.001 (ANOVA followed by Tukey’s multiple comparisons).

AQP4 polarization was calculated as the ratio of AQP4 labelling on the vessel wall to that in the parenchymal tissue directly adjacent to the vessel (Figure 6(b)). AQP4 polarization towards vessel wall was reduced significantly in peri-infarct regions of stroke compared to sham animals (F=29.46, *p*<0.001). This reduction was improved by both LOPC 8h and LOPC 24h (*p*<0.05 and *p*<0.05 respectively).

[insert Figure 6.]

## Discussion

This is the *first study*, to our knowledge, to demonstrate that exposure to low oxygen post conditioning (LOPC) is capable of reducing the accumulation and aggregation of soluble Aβ oligomers post-stroke. We have shown that a focal cortical photothrombotic stroke is responsible for inducing significant impairment of cognitive function as evaluated by the performance of the PAL task. We have demonstrated that exposure to LOPC for either 8h or 24h a day for two weeks post-occlusion was sufficient to improve the associative memory deficits induced by stroke. The improvement in cognition observed in animals exposed to LOPC was associated with marked reductions in total level of the neurotoxic protein, Aβ, both in its absolute level and in the extent of its oligomerization. LOPC was also associated with several improvements in vessel density, a reduction in vascular leakage, and restoration of AQP4 polarisation. Collectively, these findings point to the potential utility of LOPC as a therapeutic tool to improve cognition post-stroke.

In the current study, we chose to assess changes in cognition using the PAL task. The PAL test is a well-described cognitive assessment task, and was chosen as PAL deficits have been identified in patients post-stroke and in the earliest stages of Alzheimer’s disease.^29-31^ The rodent version of the PAL task used in the current study has been validated as a sensitive tool for identifying complex cognitive deficits in rodents.^18^ We identified that animals exposed to stroke exhibited robust deficits in two key metrics, namely the % of correct trials performed, which is considered to reference both learning and memory, as well as the number of repeated trials, which is considered to index perseveration.^18, 32, 33^ Both the LOPC paradigms limited these deficits to a level that was comparable to that observed in sham animals. These findings align well with other studies that illustrate the effectiveness of LOPC to improve cognitive deficits after stroke.^10, 11^

A significant focus of the work undertaken in this study was to understand the mechanisms through which exposure to LOPC improves cognition. The most obvious explanation for the pro-cognitive effects of LOPC was that the intervention is neuroprotective.^15^ Consistent with this general postulation, we observed a neuroprotective effect of LOPC using two indices. Firstly, we identified that mice exposed to LOPC for either 8h or 24h exhibited a reduced area of brain tissue loss relative to the stroke group, and secondly we demonstrated an increase in the number of NeuN+ neurons. It should be noted that glial cells, including astrocytes and microglia, have significant greater oxidative tolerance when compared to neurons. It will be of considerable interest to examine whether low oxygen exposure reduces cell death and removal or delayed injury in future studies.

An explanation for the neuronal protection conferred by LOPC was improvements within the neurovascular unit. The ability of low oxygen to stimulate growth of cerebrovasculature is well documented.^5, 6, 34^ We chose to investigate changes in vessel density using the vascular marker collagen IV, which labels the endothelial basement membrane.^22^ We identified that stroke resulted in a considerable reduction in vessel density within the peri-infarct territory, an effect that was largely attenuated via LOPC. It would be worthwhile in future studies to examine whether the new vessels are functional. Interestingly, we also identified that in addition to stimulating vessel growth, LOPC also resulted in a significant reduction in vascular leakage, as indexed by peri-vascular IgG labelling. This latter finding indicates that LOPC, in addition to stimulating vessel growth, produces a more mature vessel phenotype. Consistent with this we further observed that LOPC resulted in a restoration of AQP4 polarisation, a water channel, critical in facilitating the transit of cerebrovascular spinal fluid into the parenchyma. We also noted that LOPC resulted a mild stimulation of microglia, as evidenced by an enhanced level of Iba-1 expression. This modest enhancement is particularly interesting given the recent work demonstrating the essential role of microglia in mediating vascular repair.^35-37^ Further, characterising microglial engagement with vascular repair in the context of LOPC represents an area of future exploration.

We anticipated that if LOPC stimulated the development of the cerebrovasculature it may also promote the removal of neurotoxic proteins, such as Aβ, which is deposited at higher levels after stroke, and well recognised to disrupt neuronal function.^13, 16, 38, 39^ As expected we identified that stroked animals exhibit greater levels of the soluble oligomers of Aβ. In comparison, stroked animals exposed to LOPC exhibited significant reductions in total levels of Aβ, as well as exhibiting specific reductions in 25, 50 and 56 kDa oligomers relative to the stroke alone condition. To our knowledge, this is the first report to demonstrate that LOPC is capable of inducing robust decreases in total levels of amyloid and reducing aggregation of soluble Aβ oligomers post-stroke. It is important to recognise, however, that the relationship between the LOPC driven reduction in Aβ is associatively linked to the improvement in cognition. Future studies will be required to establish the causality of this relationship.

There are several possible explanations for the ability of LOPC to reduce the Aβ burden post-stroke. For instance, changes in Aβ may result from: changes in the rate of production of the amyloid precursor protein (APP); the rate of APP conversion into Aβ (alpha- and beta-secretase); the extent of intracellular ingestion of Aβ and the actions of degrading enzymes (NEP, ECE and IDE). Alternatively, it may be influenced by the availability of vascular bound transporters (LRP-1) and the receptor for advanced glycation end (RAGE) products involved in the excretion of Aβ or glymphatic clearance.^40-42^ While we did not quantify rates of glymphatic clearance we did identify that ability of LOPC to restore vascular polarisation of the AQP4 protein, a state which is considered to be essential for facilitating glymphatic flow.^43, 44^ We additionally observed that stroke induced disturbances in expression levels of the APP and BACE mRNA expression and that LOPC returned these to the levels observed in sham animals. We could not, however, find any evidence to suggest that LOPC altered the expression of other enzymes involved in Aβ formation or digestion including: TACE, NEP nor IDE. Modest elevations were observed in LOPC 8h exposed animals over stroke alone in mRNA expression of the transport protein LRP-1 and the RAGE products. Together these results suggest that the improvement in Aβ loading induced via LOPC is driven by a constellation of changes in the processing and removal of Aβ. However, the evidence suggests that the reduced levels of Aβ are likely the result of improved vascular density and improved vascular transportation.^45^ Further experimentation here is required, ideally using the intracranial delivery of tagged-Aβ that can be tracked in real-time.^41^

The ability of low oxygen exposure has long been recognised to be potently neuroprotective.^5^ Indeed, the therapeutic effects do not appear to be confined to the CNS, with *Nakada et al*. recently demonstrating the effectiveness of low oxygen exposure to improve heart regeneration following cardio myocyte loss.^46^ One of the principle advantages associated with reduced oxygen exposure interventions is that the approach triggers system-wide compensatory adaptions. The findings from this study further support the potential of LOPC for promoting recovery post-stroke. We have identified that exposure to a low oxygen environment for two weeks following stroke, beginning 3 days post-infarction, is sufficient enough to reduce neuronal loss, improve cognition, restore several of the vascular deficits as well se reduce the severity of the Aβ burden. Stroke is known to trigger cognitive impairment and under certain circumstance trigger the emergence of dementia-like symptoms. A therapeutic strategy to reduce Aβ and improve cognitive performance is highly desirable.^39^ As several human specific technologies already exist for controlled oxygen exposure, it is not inconceivable that future safety trials for this promising pro-cognitive intervention could be undertaken rapidly.

## Acknowledgments

We express our gratitude to HMRI Core Histology Facility for assistance with the immunohistochemistry images. We would like to acknowledge the insights and comments provided by Prof Jorgen Isgaard, Prof Neil Spratt and Dr Kirsten Coupland on early versions of the manuscript.

## Authors’ contributions

ZZ, MN and FRW designed the experiment. ZZ performed the majority of the experiments. LKO undertook all western blotting analyses and prepared the results for these data. GP undertook all mRNA analyses and prepared the results for these data. SSB, KW, MI, MK and CT assisted in the experiments. SJJ designed and prepared program for the image processing. ZZ, LKO, GP, OPO, SJJ, MN and FRW analyzed the data and interpreted the results. ZZ and FRW wrote the paper and LKO, GP, SSB, KW, MI, MK, CT, OPO, SJJ and MN revised all drafts and the manuscript.

